# CBD can be combined with additional cannabinoids for optimal seizure reduction and requires GPR55 for its anticonvulsant effects

**DOI:** 10.1101/2023.02.15.528525

**Authors:** Roshni Kollipara, Evan Langille, Cameron Tobin, Curtis R. French

**Author notes:** Corresponding Author, Curtis R. French, Associate Professor, Division of Biomedical Science, Faculty of Medicine, Memorial University of Newfoundland, St. John’s, NL, Canada A1B 5S3.

## Abstract

**Background:** Cannabis has demonstrated anticonvulsant properties, and cannabis-based medicines are approved to treat pediatric patients with severe pediatric epilepsies that are particularly refractive to approved anti-epileptic drugs (AEDs). About thirty percent of epileptic patients do not have satisfactory seizure management with AEDs and could potentially benefit from cannabis-based intervention. Here we report the use of single and combined cannabinoids to treat Pentylenetetrazol (PTZ) induced convulsions in a zebrafish model, their effect on gene expression, and a simple assay for assessing their uptake in zebrafish tissues. These data provide novel insights as to the potential of treating epilepsy with cannabinoids.

**Methods:** Zebrafish larvae were treated with cannabinoids and their seizures measured through an optimized behaviour tracking method. Cannabinoid uptake was measured with a novel HPLC-UV method. Gene expression changes were assessed using quantitative PCR (qPCR), and chemical inhibitors of potential cannabinoid receptors were used to block activity.

**Results:** Treatment with cannabinol (CBN), cannabichromene (CBC) and cannabigerol (CBG) decreased seizure intensity at lower doses than CBD when accounting for the amount of cannabinoid recovered from exposed larvae. Δ^9^-tetrahydrocannabinol (Δ^9^-THC), Δ^8^-tetrahydrocannabinol (Δ^8^-THC) were effective at higher doses. Synergistic effects were observed between CBD and other cannabinoids such as Δ^9^-THC, Δ^8^-THC, and CBG. The reduction of PTZ induced seizures via CBD is partially mediated by the G-protein coupled receptor GPR55, as pharmacological inhibition of the receptor reduced the therapeutic action of CBD. Changes in expression of endocannabinoid system (*napepld, gde1, faah, ptgs2a*) and neural (*fosab, pyya*) genes in response to phytocannabinoid treatment were observed and highlight novel mechanisms of phytocannabinoid action.

**Conclusions:** CBD can be combined with additional cannabinoids for optimal reduction of seizure activity and requires the activity of GPR55. Changes in *fosab* regulation of gene expression and endocannabinoid signalling may influence the anticonvulsant effects of cannabis, however further investigation is required.

## Introduction

Epilepsy is a complex disorder characterized by repeated seizures, which are rapid discharges of action potential in the brain that can result in uncontrolled movement and periods of lack of awareness. Epilepsy has diverse etiologies, confounded by several types of seizures and six categories of epilepsy, as per the ILAE guidelines^1^. Clinical seizure control can be difficult to achieve because approximately one third of epilepsies do not have a known cause, and greater than 30% of patients with epilepsy do not respond to currently available treatments (refractory epilepsy)^2,3^. Additionally, adverse effects to current AEDs such as drowsiness gastrointestinal distress, aseptic meningitis, and risk to the unborn often result in discontinued treatment^4^.

Animal models of epilepsy play a crucial role in epilepsy research, and some of the original AEDs were discovered after studies on animal models, as discussed by Perucca, 2019^5^. The non-competitive GABA_A_ receptor antagonist pentylenetetrazole (PTZ), for example, models spontaneous epilepsy and invokes concentration-dependent seizure-like behavior in rodents and zebrafish^6–9^. Zebrafish epilepsy models are becoming more prominent in AED candidate studies due to their high fecundity, low cost to maintain, and stereotypical behaviors that begets a high-throughput workflow. Zebrafish PTZ models have been shown to respond to current AEDs^10,11^, synthetic and naturally derived AED candidates^12,13^.

Cannabis is a versatile medicinal plant that has been used to treat convulsions since ancient times^14,15^. There are over 100 phytocannabinoids found naturally in Cannabis^16–18^. In this work we focus on 6 of the most prevalent phytocannabinoids: cannabidiol (CBD), Δ^9^-tetrahydrocannabinol (Δ^9^-THC), Δ^8^-tetrahydrocannabinol (Δ^8^-THC), cannabinol (CBN), cannabichromene (CBC), and cannabigerol (CBG), and their role in reducing seizures in an optimized zebrafish seizure assay. CBD is the active component in Epidiolex®, which the FDA and EMA have approved for treating patients 2 years and older with severe genetic childhood convulsant syndromes such as Dravet Syndrome and Lennox-Gastaut Syndrome^19–21^. Emerging evidence from animal models indicate that additional cannabinoids may also have anti-epileptic effects. CBN has been reported to display anti-epileptic effects greater than CBD in a zebrafish model of Dravet syndrome^22^. CBC is one of the most abundant cannabinoids in cannabis but has not been well studied with respect to epilepsy, but is known to exhibit neuroprotective effects in cultured mouse neural cells^23^. There is also evidence that CBC when used in combination with CBD or Δ^9^-THC can help alleviate symptoms of insomnia and depression in humans and rats^24,25^. CBG was determined to be non-psychotropic and is of interest commercially as it is abundant in some commonly used hemp plants, but little is known as to its potential anticonvulsant properties^26^.

Treatment with Δ^9^-THC in combination with CBD has been the main focus of recent studies, and has been associated with improved symptoms of neurodegenerative diseases, pain perception, and depression as reviewed by Zhang and colleagues in 2022^27^. A combination of Δ^9^-THC and CBD has also been shown to be anti-epileptic in a mouse model of Dravet Syndrome and a zebrafish model of neuro-hyperactivity^28,29^. Sole treatment with Δ^9^-THC is generally reserved for extreme medical cases because of the psychoactive effects, and can be used to treat severe emesis as shown in pilot clinical studies^30^. There is also evidence that Δ^9^-THC treatment inhibits fear learning in a zebrafish model^31^ and is being investigated for treatment of psychological illnesses such as post-traumatic stress disorder (PTSD) in humans^32^. Δ^8^-THC is a less psychoactive analog of Δ^9^-THC, and a self-reporting study on the effects of Δ^8^-THC indicated a more tolerable side-effect profile while maintaining pain relief and relaxation as compared to Δ^9^-THC^33^. There is significant value in understanding the effects of phytocannabinoids and their interactions with each other, particularly in the case of epilepsy as it has been observed in patients as well as in zebrafish models that cannabis extracts have different anti-epileptic efficacy even when normalized for dose of CBD^16^.

The mechanism by which CBD and other cannabinoids elicit their anti-epileptic properties is not well understood. While interaction with receptors of the endocannabinoid signaling system has been well documented ^34^, it is not clear which receptors are required to transduce the anticonvulsant effects of phytocannabinoids. A number of receptors have been proposed including the G-protein coupled cannabinoid receptors CBR1 and CBR2, G-protein orphan receptors such a GPF18 and GPR55, as well as transient receptor potential cation channel receptors such as TPRV1^35–39^. It is also not clear whether signaling induced by phytocannabinoids affect the synthesis of naturally endocannabinoid ligands such as anandamide (AEA) and 2-Arachidonoylglycerol (2-AG) through feedback mechanisms. Activation of CBR1 and CBR2 receptors via these endocannabinoids can inhibit neurotransmitter release^40^, and changes is 2-AG levels have been noted during seizures^41^ and potentially provide neuroprotective effects^42^. Changes in endocannabinoid synthesis could thus represent a component of the anticonvulsant mechanism of phytocannabinoids.

There are also genetic markers of seizures that may be affected by the introduction of cannabinoids that may help to elucidate their mode of action^6,10^. For example, the expression *pyya, c-fos, bdnf* are altered during PTZ induced seizures in animal models, yet it is unknown if phytocannabinoids may affect their expression as part of their therapeutic potential.

In this work, we test the use of 6 common phytocannabinoids and two known endocannabinoids for their ability to reduce seizures in zebrafish using behavioral tracking and gene expression readouts. Utilizing a novel HPLC-UV based assay to quantify cannabinoid levels in zebrafish tissues after seizure activity, we demonstrate that cannabinoids such as CBC and CBN work at lower effective doses as CBD, and that cannabinoids such as CBG may act synergistically with CBD to further reduce seizure levels. Notably, we demonstrate a clear role for the orphan receptor GPR55 in mediating the anticonvulsant effects of CBD in the zebrafish PTZ model. Incubation of larvae in the endocannabinoid 2-AG increased seizure activity and treatment with phytocannabinoids altered the expression of key genes involved in the metabolism of endocannabinoids, indicating that an effect on endocannabinoid signaling may be a key component of seizure rescue via phytocannabinoid treatment in the PTZ model.

## Methods

### Zebrafish Husbandry

Wild-type zebrafish (strain AB) were reared and staged under standard conditions as previously described^43^. All experiments were completed in accordance with Memorial University of Newfoundland’s Animal Care Committee (Approval # 20222627) and the Canadian Council on Animal Care. Zebrafish larvae were reared in embryo media, consisting of 14.97 mM NaCl, 0.50 mM KCl, 0.99 mM CaCl·2H_2_O, 0.15 mM KH_2_PO_4_, 0.05 mM Na_2_HPO_4_, 0.99 mM MgSO_4_·7H_2_O, and 0.71 mM NaHCO_3_.

### Cannabinoid Dosing and Seizure Tracking

All phytocannabinoids were purchased from Sigma Aldrich (Ontario, Canada). AEA and 2-AG were purchased from Cayman Chemicals (Michigan, USA). Larvae were collected and reared according to breeding pairs. One larva (6dpf) was added per well in a 96-well plate, and 180 μL of cannabinoid treated embryo media added. The plate was incubated at 28.5^0^C for 30 minutes, and then recorded in a tracking apparatus for 30 minutes with a 5-minute acclimation. In the dark, PTZ was added to a final concentration of 2.75 mM and the plate was recorded for 30 additional minutes after a 5-minute acclimation. Activity, measured as percent pixel change, was measured for 30 minutes using Noldus EthoVision. Seizure index is the fold change in activity, calculated by dividing the activity of a treatment by the average baseline movement (activity prior to PTZ exposure).

### Phytocannabinoid Extraction

Visual description of the protocol can be found in Figure 1. Larvae were pooled in groups of 12 according to treatment received, rinsed twice in ice cold embryo media, and euthanized by incubation on ice. Excess embryo media was removed, and samples stored at −80°C until extraction. All steps of extraction were completed at 4-8°C. 200 μL of extraction solvent (methanol with 0.1% formic acid) was added to the larvae, which were then bead beaten for 3 minutes in an equal volume of 1 mm glass beads. Homogenate was then centrifuged for 30 minutes at 18,000 x G and the supernatant 0.2 μm filtered prior to HPLC analysis.

**Figure 1.**
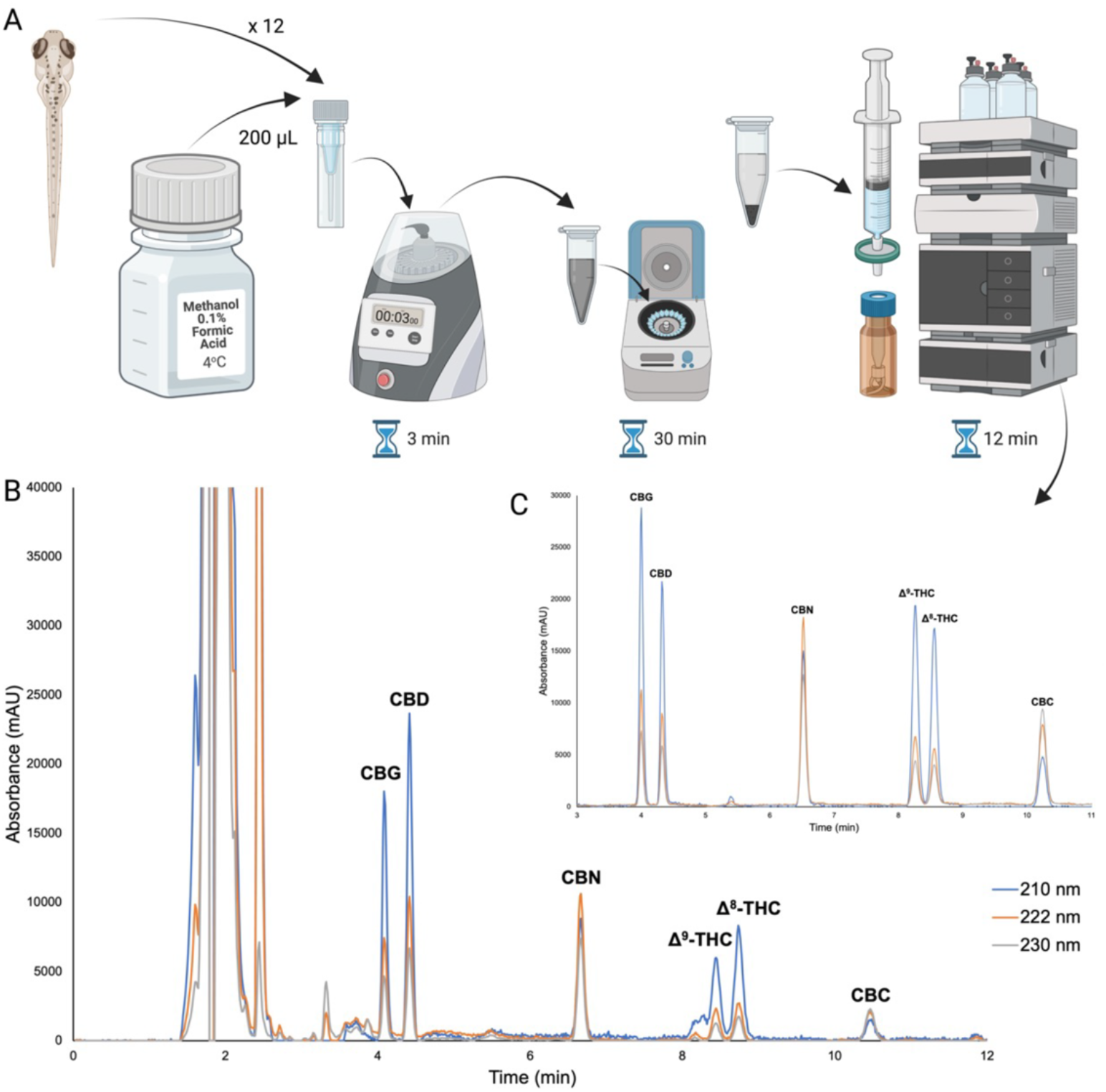
Overview of the cannabinoid extraction and HPLC method presented. A) Graphical summary of cannabinoid extraction protocol from zebrafish. B) Representative chromatogram of extraction of 12 pooled larvae treated with 6 cannabinoids at 4 μM. C) Representative chromatogram of 2.5 μg·mL^-1^ mixed standard of cannabinoids in extraction solvent. The three traces depict absorption at different wavelengths as stated in the legend. The wavelength with the highest relative absorbance was used for quantification of standards and samples. Peaks are labelled with corresponding compound abbreviations.

### HPLC Analysis of Cannabinoids

Analytical HPLC was carried out on an HP 1050 system equipped with two HALO 2.7 μm C18 100 x 4.6 mm columns (200 mm total length). A diode array detector monitored 200-400 nm. Extracted wavelength chromatograms at 210 nm for CBG, CBD, Δ^9^-THC, and Δ^8^-THC, 222nm for CBN and 230 nm for CBC were used for quantification using TargetLynx software (Waters Corp, Mass, USA), and can be found in Figure 1. Mobile phase consisted of an isocratic mixture of 17.5:82.5 water 0.1% formic acid: acetonitrile 0.1% formic acid, v/v at a flow rate of 1 mL min^-1^ resulting in a system backpressure of approximately 20 MPa. Triplicate 20 μL injections were performed per sample. External standards of the cannabinoids in extraction solvent were used to generate calibration curves (0, 0.1, 0.25, 0.5, 1, 2.5, 5 and 10 μg mL^-1^).

### RT-qPCR

Larvae (6 dpf) were treated using the same method described for seizure tracking and pooled in groups of 12 for RNA isolation. RNA was isolated using the PureLink™ RNA Mini Kit (Invitrogen) with on-column DNAase treatment using the PureLink™ DNase Set (Invitrogen) and cDNA synthesized using the High-Capacity cDNA Reverse Transcription Kit (ThermoFisher). qPCR was performed with Taqman Gene Expression Master Mix and Taqman assays (FAM) for *fosab* (Dr03100809_g1),*pyya* (Dr03138152_m1), *gde1* (Dr03434842_m1), *napepld* (Dr03117925_m1), *faah* (Dr03093136_m1), *ptgs2a* (Dr03080325_m1), and *ptgs1* (Dr03087197_m1) using a ViiA 7 qPCR machine. Wells were duplexed with the endogenous control, TATA-box binding protein (*tbp*) (VIC). Data was analyzed using the ΔΔCT method and presented as means ± standard error of the mean. Two biological replicates of 12 pooled larvae, with three technical replicates for each sample, were analyzed.

### Statistics

Significance was determined by one-way ANOVAs and post-hoc Tukey HSD using astatsa.com (copyright Navendu Vasavada). Simple statistical calculations such as mean, standard deviation, and standard error of the mean were completed on Microsoft Excel.

## Results

### Development of HPLC method

Phytocannabinoids are highly related compounds that have similar chemical properties, therefore optimization steps with analytical standards were necessary prior to processing tissue samples. A rapid and simple extraction of cannabinoids from pooled larvae is presented (Figure 1). The analytical figures of merit for the presented method are in Table S1. CBG eluted at 4.0 min, followed by CBD at 4.3 min, CBN at 6.6 min, Δ^9^-THC at 8.3 min, Δ^8^-THC at 8.6 min, CBC at 10.3 min. The lower limits of detection and quantification (LLOD and LLOQ, respectively) are within the useful limits for dosing larval zebrafish with cannabinoids (Table S1). Upper limits of quantification are well above the practical dosing range for larval zebrafish. Inter-day variability of the method was determined through injection of a 1 μg mL^-1^ mixed standard on ten different days (n=10), giving an average percent deviation of 5.3% with individual cannabinoids ranging from 1.3 to 8.6% (Table S1).

Calibration curves were created for each of the six cannabinoids in this study using the developed HPLC method (Figure S1). The procedure for extraction of cannabinoids from zebrafish larvae partitions cannabinoids into the methanol supernatant, leaving insoluble biomatter as a precipitate. To ensure complete extraction of cannabinoids from larvae, we conducted three sequential extractions on samples of treated larvae and found that a single extraction, on average for all compounds, recovered 95.7% while the second and third extractions yielded 2.4% and 1.8%, respectively. Individual recoveries from single extractions ranged from 92.0% to 98.5% (Figure S2).

### Individual treatments of cannabinoids

Zebrafish larvae were first given treatment with individual cannabinoids at physiologically relevant concentrations, with the intention to compare results to the established anti-epileptic CBD (Figure 2). Seizure activity was measured as seizure index, which is a fold change in activity, and larvae were then analyzed for cannabinoid tissue accumulation using HPLC. All cannabinoids showed a reduction in seizure activity at a 4 μM treatment (Figure 2A-F). Accumulation of cannabinoid in larvae at the lowest dose that caused a statistically significant reduction in seizures is as follows: 1.6 ± 0.1 ng/larva CBN, 1.7 ± 0.3 ng/larva CBC,, 11.9 ± 1.1 ng/larva CBD, 15.0 ± 2.5 ng/larva CBG, 30.6 ± 1.1 ng/larva Δ^8^-THC, and 48.3 ± 3.3 ng/larva Δ^9^-THC.

**Figure 2.**
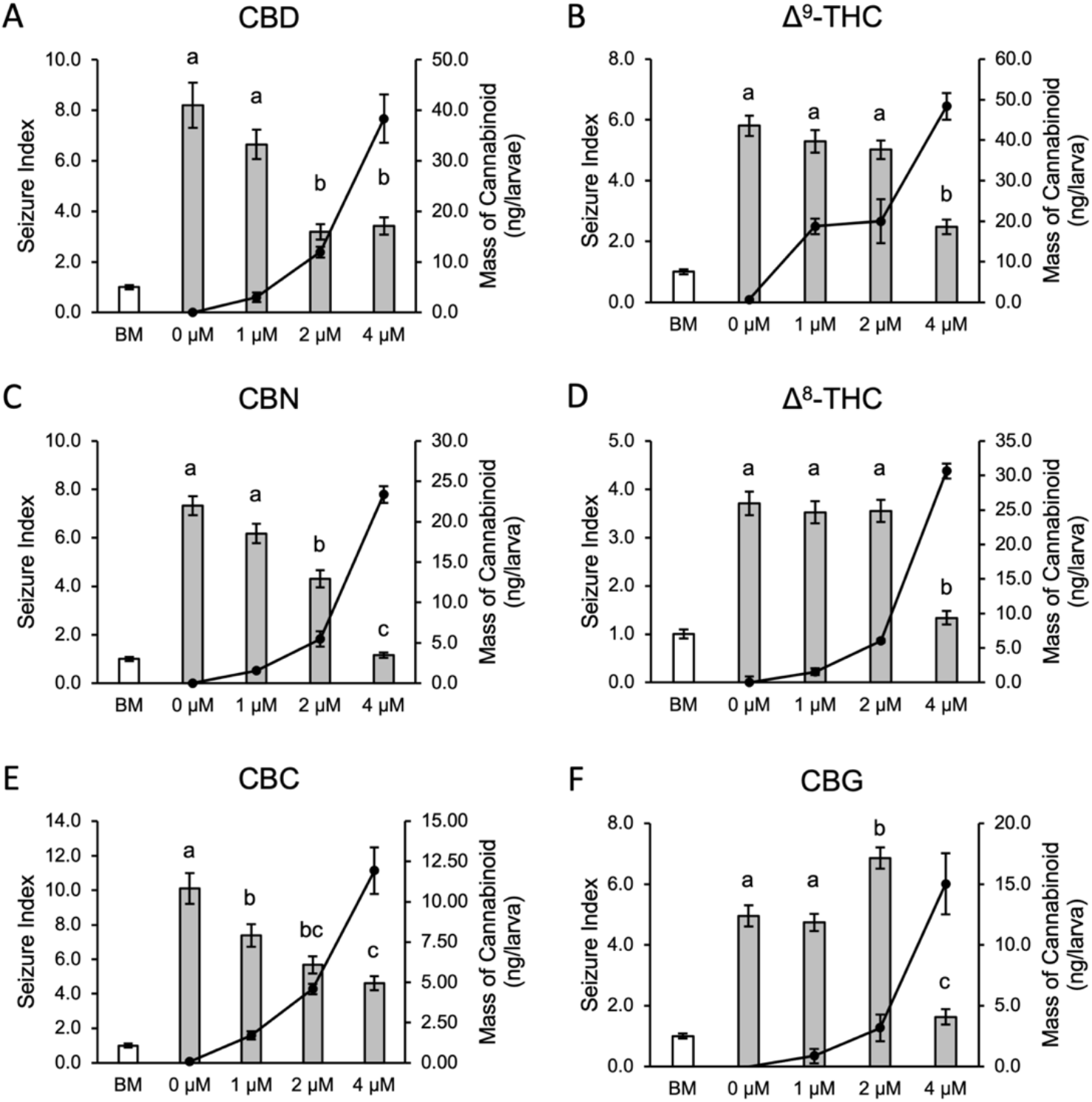
A-F: Seizure reduction and cannabinoid concentration following individual dosing of cannabinoids. Seizure Index is fold change in movement, displayed as bars (error bars are SEM, significance calculated by one-way ANOVA, alpha<0.05). BM: baseline movement before addition of PTZ. Grey bars are movement after addition of PTZ to final concentration of 2.75 mM, with x-axis labels corresponding to concentration of cannabinoid. “0 μM” treatment is the vehicle control. n_BM_ = 288. A: n_treatment_ = 71-72, B: n_treatment_ = 71-72, C: n_treatment_ = 68-72, D: n_treatment_ = 71-72, E: n_treatment_ = 71, F: n_treatment_ = 70-72. HPLC quantification of mass of cannabinoid per larva is shown as connected points (error bars are SD), n=8 samples per data point with triplicate injection.

As reported previously, CBD showed seizure relief at 2μM (61% reduction p<0.01) and 4 μM treatments (58% p<0.01, Figure 2A). This was also observed in individual treatments of CBN where 2 μM reduced seizure index by 41 % (p<0.01) and 4 μM treatment reduced seizure index by 84 % P< 0.01, Figure 2C). Similar reductions of seizure index were observed for and CBC, where 2 μM reduced seizure index by 44 % (p<0.01) and 4 μM treatment reduced seizure index by 54 % (p< 0.01), however there was also a statistically significant reduction in seizures at a 1 μM CBC treatment (27 % p<0.05, Figure 2E). Accumulation of CBD, CBN, and CBC in tissue decreased respectively, with CBC observed to have the least accumulation in tissues with the doses tested in this study. CBC and CBN caused a statistically significant reduction in seizures with the lowest body concentration compared to the other cannabinoids in this study. The accumulation of Δ^8^-THC in larval tissue showed an expected dose-dependent increase (Figure 2D). A plateau was observed in Δ^9^-THC accumulation, where a difference in accumulation was not seen between 1 and 2 μM treatments but was observed after 4 μM treatment (Figure 2B). A higher mass of Δ^9^-THC accumulated in tissue compared to Δ^8^-THC. CBG only provided seizure relief at a dose of 4μM, where a 67 % reduction in seizure index was observed p< 0.01) but interestingly there was a 38% increase in seizure index with 2 μM treatment (p <0.01 Figure 2F).

### Mixed treatments of cannabinoids

The anti-epileptic properties of CBD have been well documented, with some reports of an enhancement of these properties after a combined treatment with other cannabinoids^16,28^. CBD was paired with each of the five remaining cannabinoids in this study, with the results compared to individual treatments (Figure 3). As previously reported^28^, increased anti-epileptic effects are observed by combining CBD with Δ^9^-THC (Figure 3A), which resulted in a 91% reduction in seizure index (p<0.01). Increased anti-epileptic effects were also observed in treatments where CBD was paired with CBG (Figure 3B), whereby combined treatment resulted in a 95% reduction in seizure index (p< 0.01). The seizure reduction from combined CBD and Δ^8^-THC treatment reached 92% (Figure 3C, p<0.01). Paired treatments of CBD with CBN or CBC did not reduce seizure index more than each individual treatment (Figure 3D and E).

**Figure 3.**
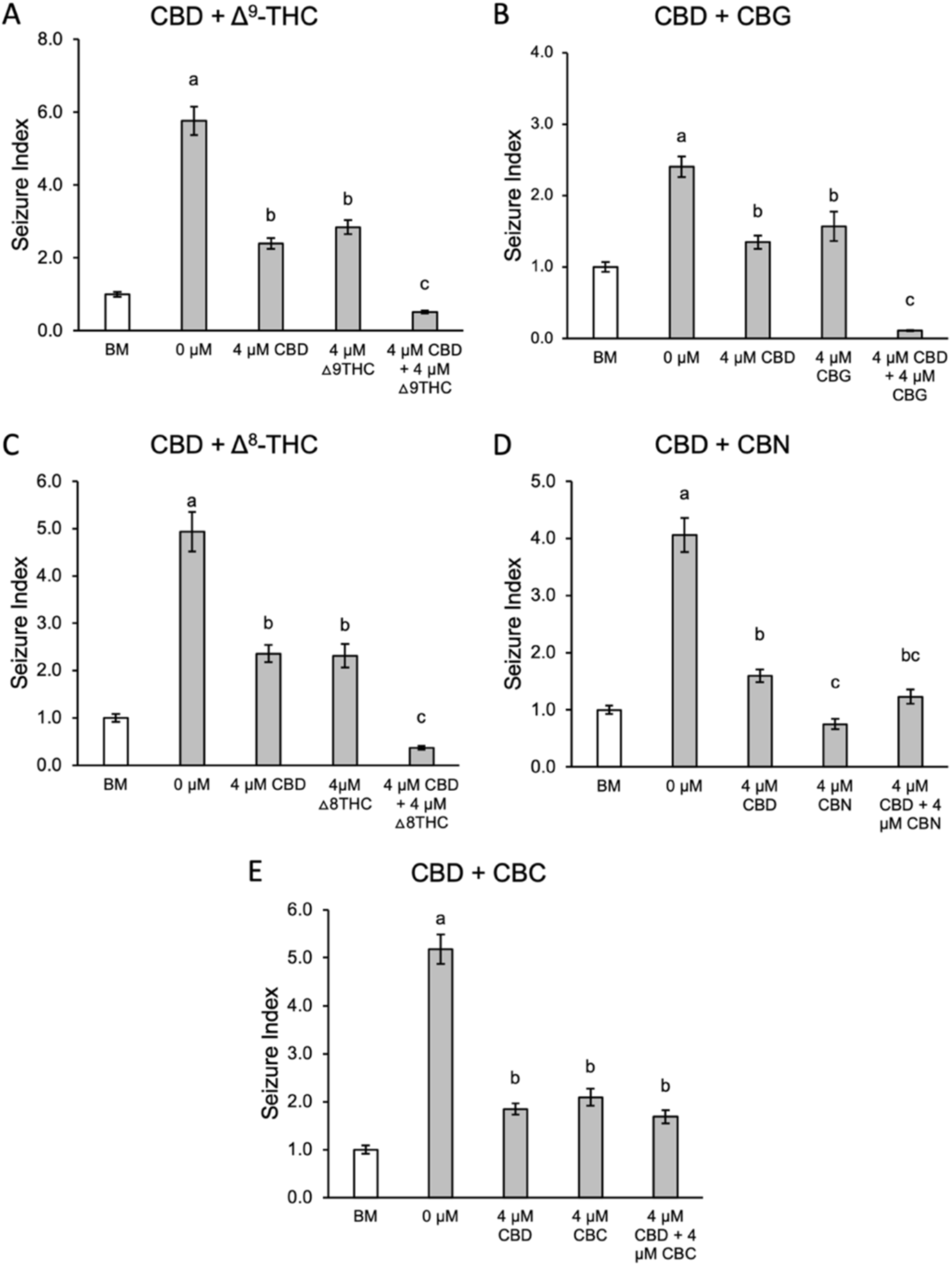
A-E: Seizure tracking comparing the effects of dosing with individual verses paired cannabinoids. Seizure Index is fold change in movement, error bars are SEM. BM: baseline movement before addition of PTZ. Grey bars are movement after addition of PTZ to final concentration of 2.75 mM, with x-axis labels corresponding to dose of cannabinoid. “0 μM” treatment is the vehicle control. Significance is noted by data labels (one-way ANOVA with Tukey HSD, p<0.05); bars with a different letter data label are significantly different. n_treatment_ = 72, n_BM_ = 288.

### GPR55 mediates CBD seizure relief

CBD treatment was paired with antagonists of receptors that likely mediate seizure relief (Figure 4). CB1R is antagonized by AM251^44^, GPR55 is antagonized by ML-193^45^, and GPR18 is antagonized by PSB-CB5^46^. Whereas seizure relief from CBD treatment reached 57% in this assay, blocking GPR55 before CBD treatment resulted in only a 30% reduction in seizure index (p<0.05), however seizures were still reduced as compared to the vehicle control. The anti-epileptic effect of CBD was not affected by blocking of CB1R or GPR18.

**Figure 4.**
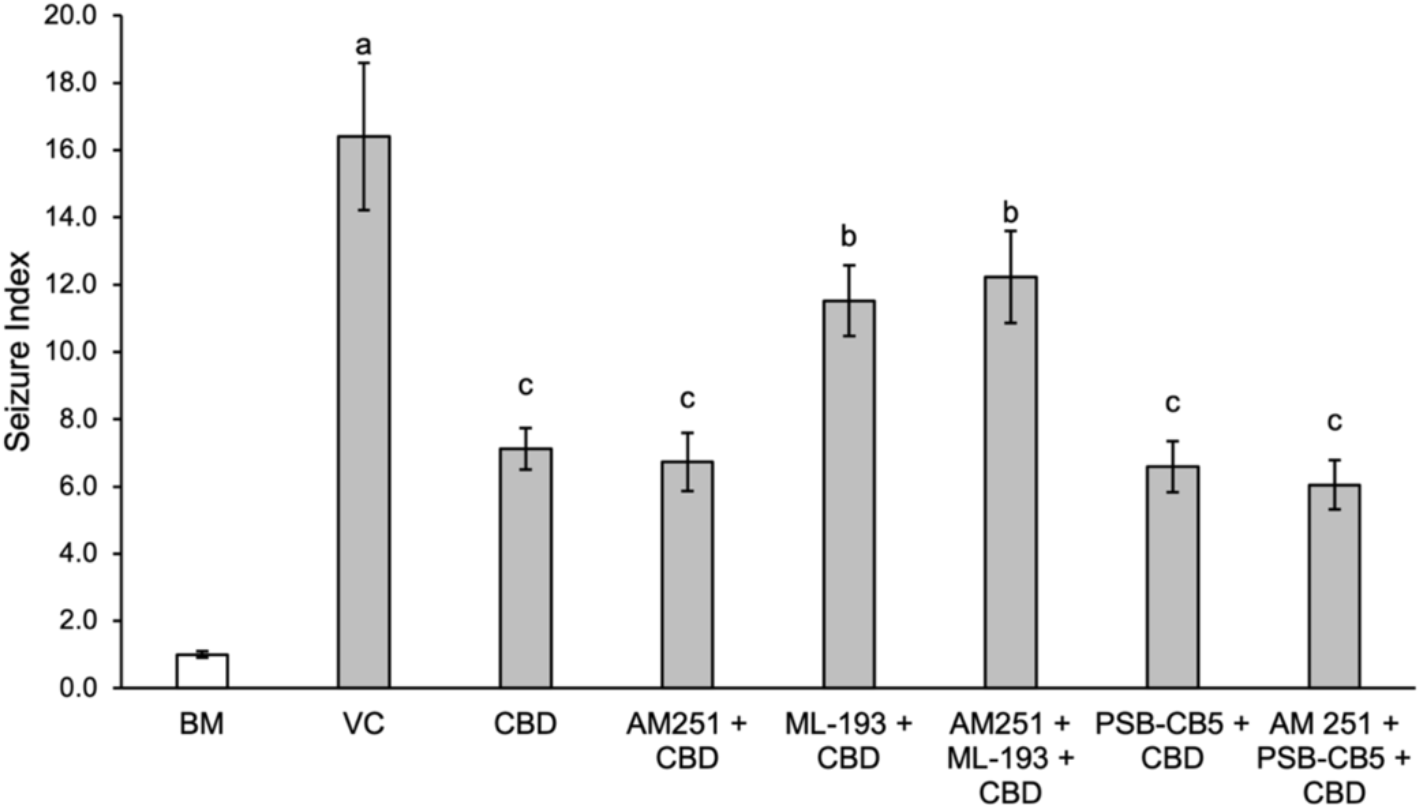
Seizure behavior after treatment with CBD and receptor antagonists. BM: baseline movement before addition of PTZ; Grey bars are movement after addition of PTZ to final concentration of 2.75 mM, with x-axis labels corresponding to treatment. VC: vehicle control. Significance is noted by data labels (oneway ANOVA with Tukey HSD, p<0.05); bars with a different letter data label are significantly different. n_treatment_ = 34-36, n_BM_ = 288.

### Endocannabinoids and seizure activity

Given the proposed overlap in the mechanism of function between endocannabinoids and phytocannabinoids, the potential of treating epilepsy with endocannabinoids was investigated (Figure 5). Larvae were treated using the same protocol as individual phytocannabinoid treatments and increase in seizure index was observed at when larvae were incubated in 1 μM 2-AG, but not at doses of 2 and 4 μM. There was also no difference in seizure activity due to incubation of larvae in 1-4 μM AEA and it was determined that there is no seizure relief produced by treatment with 2-AG or AEA when applied at concentrations ranging from 1-4 μM.

**Figure 5.**
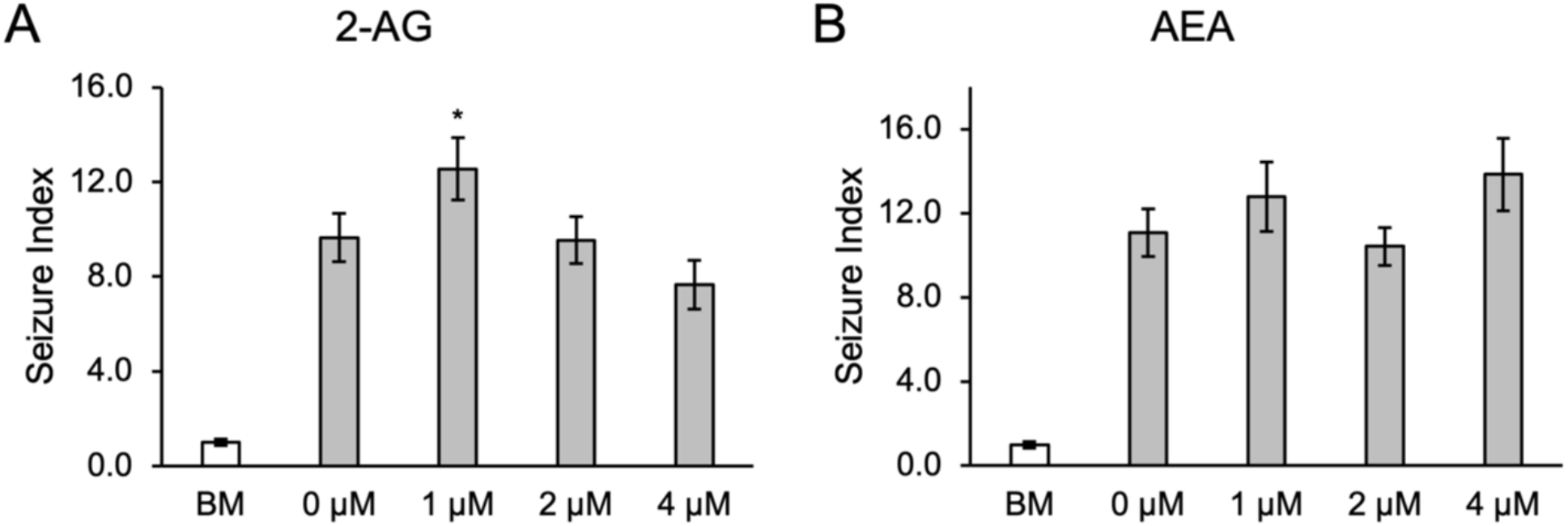
Dose-response of epilepsy model to endocannabinoid treatment as per the chart titles. BM: baseline movement before addition of PTZ. Grey bars are movement after addition of PTZ to final concentration of 2.75 mM, with x-axis labels corresponding to dose of endocannabinoids. “0 μM” treatment is the vehicle control. Error bars are SEM, significance calculated by one-way ANOVA, alpha<0.05. A: n_BM_ = 192, n_treatment_ = 47, 46, 44, 43 in the order presented in the graph. B: n_BM_ = 160, n_treatment_ = 40.

### Cannabinoid induced changes in expression

Based on observed anti-epileptic properties, changes in expression of neural and endocannabinoid pathway genes were evaluated in our epilepsy model after treatment with CBD, CBN, and Δ^9^-THC (Figure 6). Using qPCR on whole body RNA, seven gene targets were investigated: *fosab, pyya, napepld, faah, gde1, ptgs2a*, and *ptgs1*. Expression of *fosab* was increased 3.3 fold (p< 0.01) via treatment with PTZ as previously shown^8^ and was further increased when combined with cannabinoid treatments. Pre-incubation of PTZ treated embryos with CBD resulted in a 9.9± 0.5 fold induction (p< 0.01), pre-incubation in CBN resulted in 6.5 ± 0.3 fold induction (, p< 0.01), and Δ^9^-THC resulted in a 7.7 ± 0.4 fold induction (p< 0.01). While PTZ exposure did not statistically affect the expression of *pyya*, it was significantly increased 1.4 ± 0.1 fold with combined PTZ and CBD treatment when compared to wild type siblings (p< 0.05). Expression of *napepld* was reduced in combined PTZ/ Δ^9^-THC treated embryos compared to PTZ treatment alone (0.8 ± 0.05 fold p< 0.05). Similarly, Δ^9^-THC was the only treatment to significantly decrease expression of *gde1* (0.8 ± 0.1 fold, p< 0.05), when combined with PTZ. Expression of *ptgs2a* was unchanged after treatment with PTZ alone but increased 1.8 ± 0.1 fold with CBD and PTZ treatment (p< 0.01) and 1.6 ± 0.2 fold for combined PTZ and CBN treatment (p< 0.05). Expression of *ptgs1* was not significantly different in any treatment as compared to wildtype expression. Expression of *faah* was increased after treatment with Δ^9^-THC as compared to the PTZ treated embryos (1.5 ± 0.2 fold p,0.01), but not compared to WT siblings.

**Figure 6.**
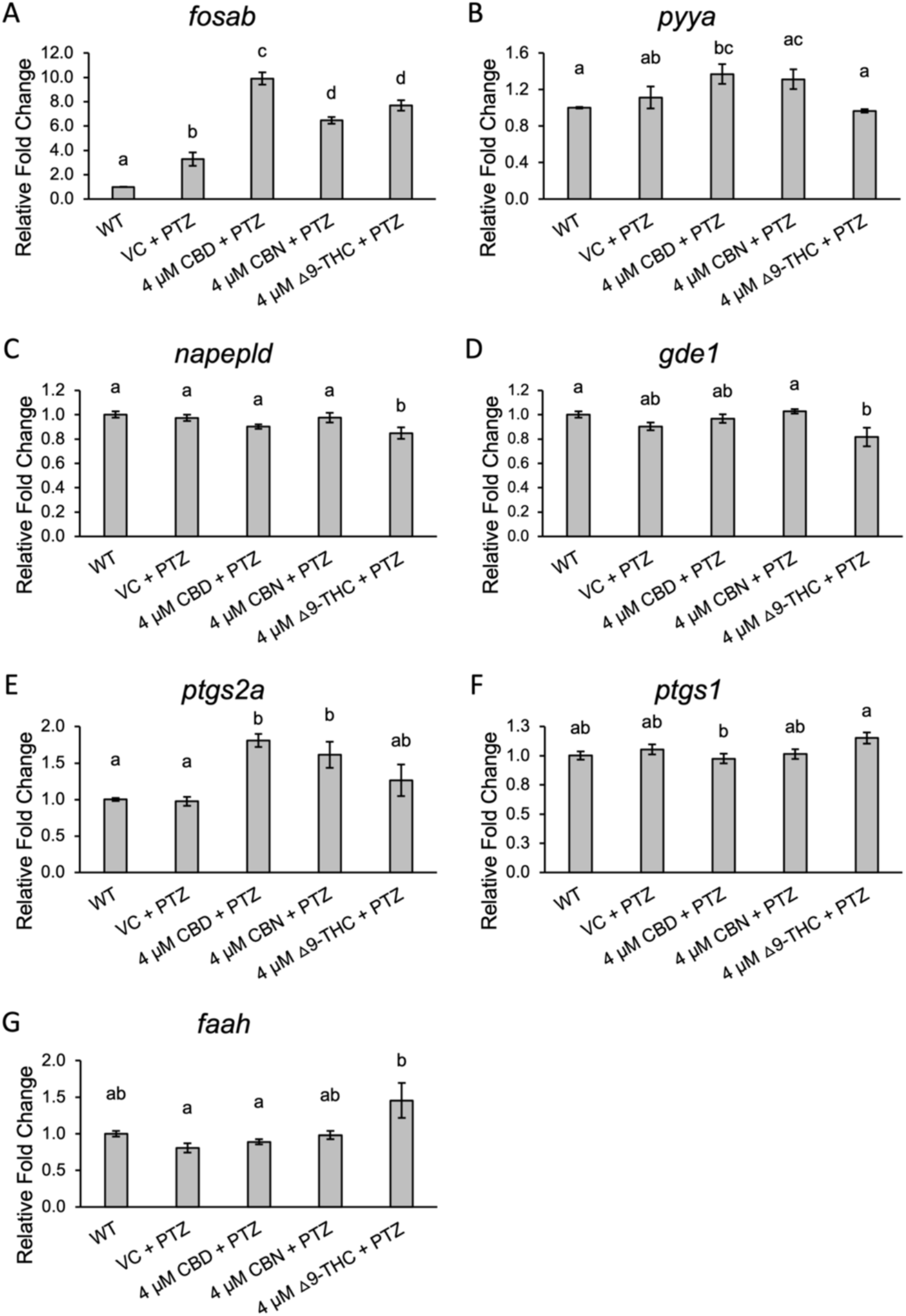
qPCR expression analysis of seizure markers and endocannabinoid system genes. Methanol was used as the vehicle control, with 4 μM cannabinoid doses. Significance is noted by data labels (one-way ANOVA with Tukey HSD, p<0.05); bars with a different letter data label are significantly different. Error bars are SEM.

## Discussion

Analysis of experimental samples with phytocannabinoids displayed potent anticonvulsant effects with increasing amount of cannabinoid quantified per larva at each increasing dose given. The use of a novel HPLC method allowing for direct quantitation of cannabinoids in larvae builds upon previous studies^47,48^, allowing for the analysis of anticonvulsant activity per mass of cannabinoid that accumulates in larval tissues. The method runtime of 12 minutes is suitable for medium-throughput analysis, with separation of 6 cannabinoids.

Our seizure tracking data were consistent with a breadth of previous literature supporting the use of CBD as one of the most effective cannabinoids in the reduction of seizures. CBN and CBC also showed a potent reduction of seizures at 1 μM dosage and interestingly, accumulated in tissue with similar values of 1.6 ± 0.1 ng/larva for CBN and 1.7 ± 0.3 ng/larva for CBC, the lowest abundance quantified that was associated with a reduction in seizures. Though previously reported to possess anti-epileptic properties, Δ^9^-THC^27–29^, and its structural isomer Δ^8^-THC, only provided seizure rescue at the highest concentrations used in this study (4 μM) and required higher accumulation within larvae for such effects. Interestingly, a 2 μM dose of CBG repeatedly increased seizure index significantly to a level higher than the vehicle control, yet higher doses reduce seizure index. CBG has a broad range of binding affinities for a unique range of receptors, which has been heavily reported to change based on concentration^49^, which is likely the reason behind this unexpected dose-response trend.

We evaluated seizure behaviors after treatments of CBD paired with each of the remaining 5 cannabinoids in this study. Strikingly, we observed synergistic effects between CBD and cannabinoids that when dosed individually were only effective at 4 μM, namely Δ^9^-THC, Δ^8^-THC, and CBG. These data supports previously reported observations that CBD treatment can provide more effective seizure reduction when combined with other cannabis compounds such as Δ^9^-THC^28^, however to the best of our knowledge this is the first report of increase seizure relief from combined CBD + CBG and CBD + Δ^8^-THC. The synergistic effect of CBD + CBG treatment is of clinical relevance as it could potentially provide improved seizure relief, but further study is needed as overall activity with this combined treatment was below baseline and could thus indicate sedation as opposed to true seizure relief. CBN and CBC, which show antiepileptic effects when used individually, did not show any increased in seizure relief when given together with CBD. This could indicate overlapping mechanisms of action, with saturation via either compound at 4 μM dosage. While a number of studies have utilized chemically characterized cannabis plant extracts that contain multiple cannabinoids and other potentially clinically relevant molecules^16,50^, to the best of our knowledge this is the first report of systematic paired dosing of CBD with five other cannabinoids in an epilepsy model. The use of single anticonvulsant cannabinoids such as CBN and CBC may provide an avenue for treatment with different side effect profiles than CBD. Moreover, as tolerance to treatment with CBD and Δ^9^-THC have been reported^51,52^, using additional cannabinoids or combining with CBG for increased effect may offer avenues for avoiding resistance to treatment over time. The combined use of CBD and CBG may offer a treatment option for increased relief similar to the observed effects of combined CBD and Δ^9^-THC use, but without the psychoactive effects of Δ^9^-THC.

After establishing the effects of treatment with individual and combined cannabinoid treatments, we sought to further investigate the mechanism of action of CBD mediated seizure relief. By treating larvae with CBD as well as antagonists of CB1R, GPR55, and GPR18, a potential mechanism of action was determined by observing which conditions negatively affected seizure relief. The data shows that CBD acts through GPR55 as one modality to reduce seizures. When GPR55 is blocked, CBD induced seizure relief is significantly reduced but not eliminated, suggesting there is more than one mechanism of action. When both GPR55 and CB1R are blocked, the reduction in seizure relief is slightly more than inhibition of GPR55 alone, but this was not statistically significant. This suggests that CB1R may play a minor role in mediating CBD’s anti-epileptic effects that is supported by reports of CBD having low affinity to CB1R, however further investigation is required. While a number of studies have suggested that cannabinoids can bind to GRP18^53,54^, blocking GPR18 showed no effect on CBD induced seizure reduction and thus this cannabinoid receptor likely does not transduce the anticonvulsant effects of CBD.

Given the role of the endocannabinoid receptors in mediating the anticonvulsant effects of phytocannabinoids, we sought to better understand the potential role of endocannabinoid ligands in the production of, or relief from seizures. Endocannabinoid ligands and their resulting signaling pathways generally considered protective with respect to seizures^41,55^, however there are conflicting data. For example, while AEA has been shown to have anticonvulsant effects, loss of the enzyme FAAH that catabolizes AEA to arachidonic acid (AA) renders AEA treatment proconvulsive^56,57^. This suggests that AA may actually mediate the anticonvulsant effects observed from AEA treatment, or that the ratio of AEA to AA may be important in regulating neuronal hyperexcitability. As changes in endocannabinoid ligand levels have been noted in cannabis users, we hypothesized that the induction of seizures, or their relief through phytocannabinoid exposure, may alter endocannabinoid ligand synthesis. The expression of *napepld, gde1* (synthesis of AEA) and *faah* (breakdown of AEA) was not significantly affected by PTZ treatment, which suggests that they are not involved in seizure induction and progression in this model. There was however a significant reduction of *napepld* and *gde1* expression after Δ9-THC exposure in PTZ treated embryos. AEA and Δ^9^-THC have been reported to act on the same receptors^27,34,58^ and this further supports literature on the putative mechanism of action of Δ^9^-THC. Expression of *faah*, required or the breakdown of AEA to AA, was significantly increased in PTZ exposed embryos pretreated with Δ^9^-THC when compared to PTZ treatment only. Together, these results suggest that Δ^9^-THC may change the availability of AEA in PTZ seizure models, or alter the ratio of AEA to AA, however this needs to be tested directly.

Expression of the zebrafish ortholog of *cox-2 (ptgs2a*) was increased after cannabinoid treatment when compared to wildtype and PTZ treated embryos, with results from CBD and CBN treatment being statistically significant. COX-2 is responsible for the oxidation of a minor amount of 2-AG and AEA^34,59^, and thus endocannabinoid levels may be reduced after CBD or CBN treatment, although this was not directly tested. These data indicate that a feedback system which may negatively regulate endocannabinoid synthesis could be triggered by certain phytocannabinoids. As we have demonstrated that incubation of larvae in endocannabinoids can increase seizures in our model, a reduction of endocannabinoid synthesis could potentially represent a component of the anticonvulsant effects of Δ^9^-THC, CBD, and CBN.

Lastly, we assessed known neuronal markers of seizure activity to determine whether their alteration by phytocannabinoids represent a path toward seizure relief. The neuronal activity marker c-Fos (*fosab*) commonly serves as a marker for seizure progression in epilepsy studies^6,60^. Some AEDs have been shown to decrease *fosab* expression to wild-type levels^10^. While incubation of larvae in PTZ more than doubled the expression of *fosab*, we found that treatments of CBD, CBN, and Δ^9^-THC dramatically increased *fosab* expression further, even though larvae in these treatment groups displayed reduced epileptic behavior. Treatment with CBD or Δ^9^-THC was reported to cause a dose-dependent increase in *fosab* expression in 96 hpf embryos^48^, and a fear-learning study found that only specific brain structures increase *fosab* expression following Δ^9^-THC treatment^31^. While these data do not support a model *fosab* as a marker of seizure progression, they do support a hypothesis that larvae may increase expression of *fosab* after seizures, and the further increase with phytocannabinoid treatment may provide therapeutic benefit.

Previous work has demonstrated that seizures induced expression the signaling peptide encoded by *pyya*^10,61^. In our study, there was no statistically significant increase of *pyya* expression after PTZ treatment, however PTZ dose and the age of larvae used differ between studies. The expression of *pyya* is increased in larvae treated with CBD subsequent to PTZ exposure compared to wild type larvae, indicating that CBD and PTZ may be activating similar pathways toward *pyya* regulation. Some studies have associated peptide YY with the inhibition of seizures and neuroprotection from stress^62,63^, and thus its increase after combined CBD and PTZ exposure, but with PTZ alone, may indicate a compensatory mechanism for seizure relief.

In conclusion, a robust and high-throughput behavioral analysis method yielded sensitive and accurate measurements of seizure reduction resulting from phytocannabinoid treatment. A novel HPLC method was employed for the sensitive and reliable measurement of six therapeutically important cannabinoids in pooled zebrafish larvae. When dosed individually, CBC and CBN were observed to have significant anti-epileptic effects at lower doses than what was observed with CBD and accumulated in tissues at lower levels indicating that may act as more potent therapeutics. CBD induced seizure relief is partially mediated by GPR55, while additional cannabinoids can be used in conjunction with CBD for increased therapeutic value. The increased expression of *fosab* during seizures is further increased with phytocannabinoid treatment, suggesting that increasing *fosab* expression may be a compensatory mechanism. While the expression of other neural makers was affected by phytocannabinoid treatment, there lack of expression changes during PTZ exposure suggests they may not be involved in the mechanism of seizure development but represent alternative pathways that may be manipulated for therapeutic benefit. The role of endocannabinoid system in phytocannabinoid seizure relief, including genes involved in the synthesis and breakdown of AEA and 2-AG and the GPR55 receptor, are supported by this study.

## Acknowledgements

This research was funded through a research contract with CEPG Consulting and Design Inc, and conducted under their research and development license from Health Canada (LIC-Z2ERZBAS9U-2020). Funding was also received from Epilepsy NL, and in the form of a Mitacs Accelerate Fellowship. R.K. and E.L. both received SGS baseline funding from the School of Graduate Studies, Memorial University of Newfoundland. C.T. received the Undergraduate Student Research Award from the National Sciences and Engineering Research Council. We would like to thank Dr. Matthew Rise (Memorial University of Newfoundland and Labrador) for the use of his behavioral tracking system and software.

## Author contributions

R.K. and C.F. conceived the experiments. R.K., E.L., and C.T. conducted the experiment(s), and R.K. performed statistical analysis and figure generation. All authors reviewed the manuscript.

