## Supplemental table and Figures for "CBD can be combined with additional cannabinoids for optimal seizure reduction and requires GPR55 for its anticonvulsant effects"

**Supplementary Information**

**Table S1**. Cannabinoid retention times, quantification wavelengths and analytical figures of merit of the presented method. RT = retention time; Quant. WL = quantification wavelength; LLOD = lower limit of detection; LLOQ = lower limit of quantification; IDV = inter-day variability.

| **Analyte** | **RT (min)** | **Quant. WL (nm)** | **LLOD (µg⋅mL^-1^ for 20 µL inj.)** | **LLOQ (µg⋅mL^-1^ for 20 µL inj.)** | **LLOD**  **ng/larva** | **LLOQ**  **ng/larva** | **Linear range and R^2^, n=3** | **IDV**  **(%, n=10)** |
| --- | --- | --- | --- | --- | --- | --- | --- | --- |
| CBG | 4.0 | 210 | 0.02 | 0.07 | 0.27 | 1.01 | 0.1-10  0.9999 | 5.6 |
| CBD | 4.3 | 210 | 0.02 | 0.08 | 0.28 | 1.13 | 0.1-5  0.9997 | 7.7 |
| CBN | 6.6 | 222 | 0.01 | 0.05 | 0.17 | 0.66 | 0.1-10  1.00 | 3.1 |
| Δ^9^-THC | 8.3 | 210 | 0.01 | 0.08 | 0.12 | 1.04 | 0.1-5  0.9997 | 5.9 |
| Δ^8^-THC | 8.6 | 210 | 0.01 | 0.10 | 0.17 | 1.45 | 0.1-5  0.9994 | 8.6 |
| CBC | 10.3 | 230 | 0.03 | 0.08 | 0.35 | 1.11 | 0.1-10  0.9999 | 1.3 |

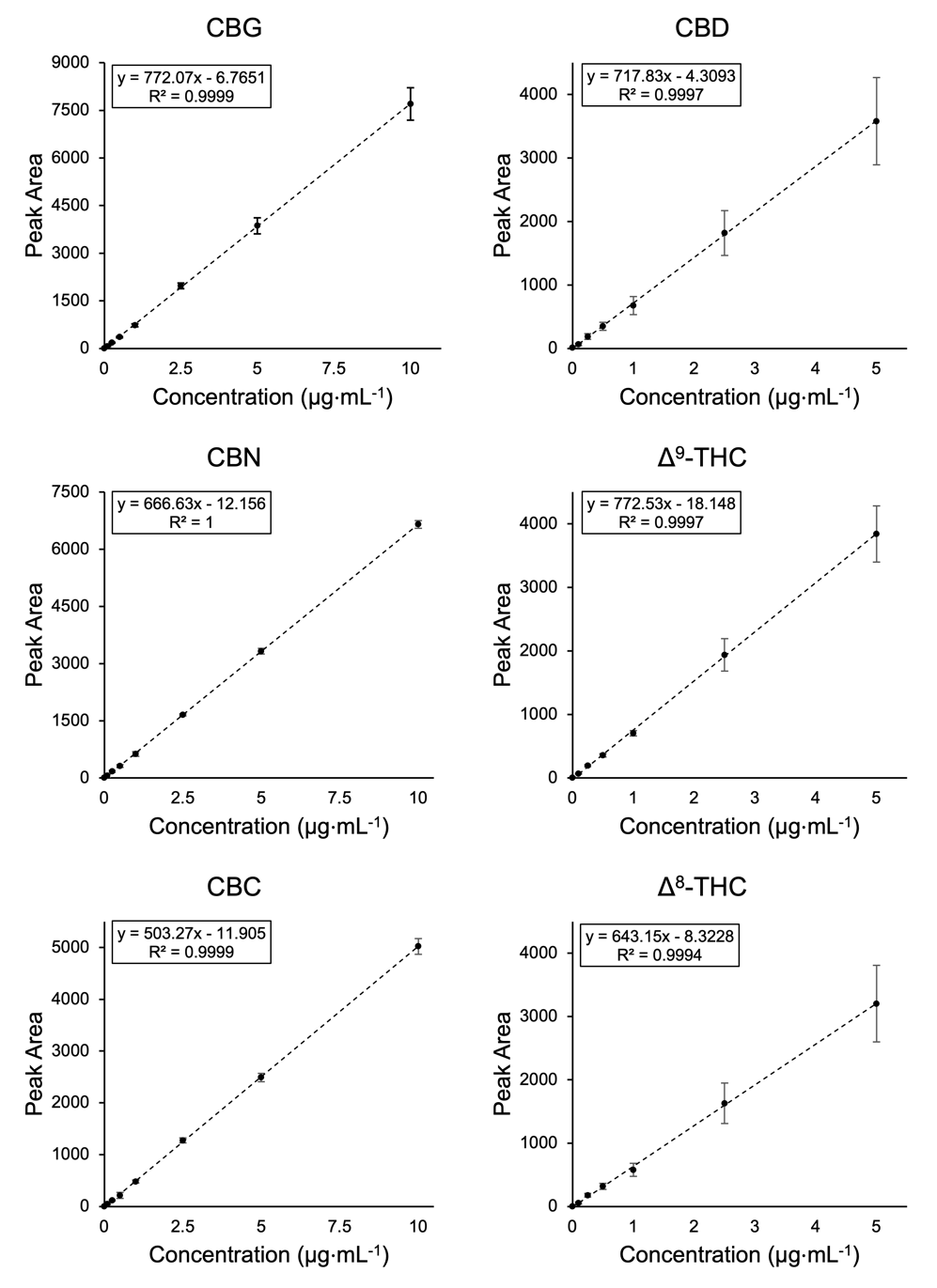

**Figure S1. External calibration curves across the linear range of each compound, presented as average of three different days.** Error bars represent the standard deviation between the days. Linear regression analysis is plotted as dotted lines with the equation for the line of best fit and the linear correlation coefficient (R^2^) in the overlaid boxes.

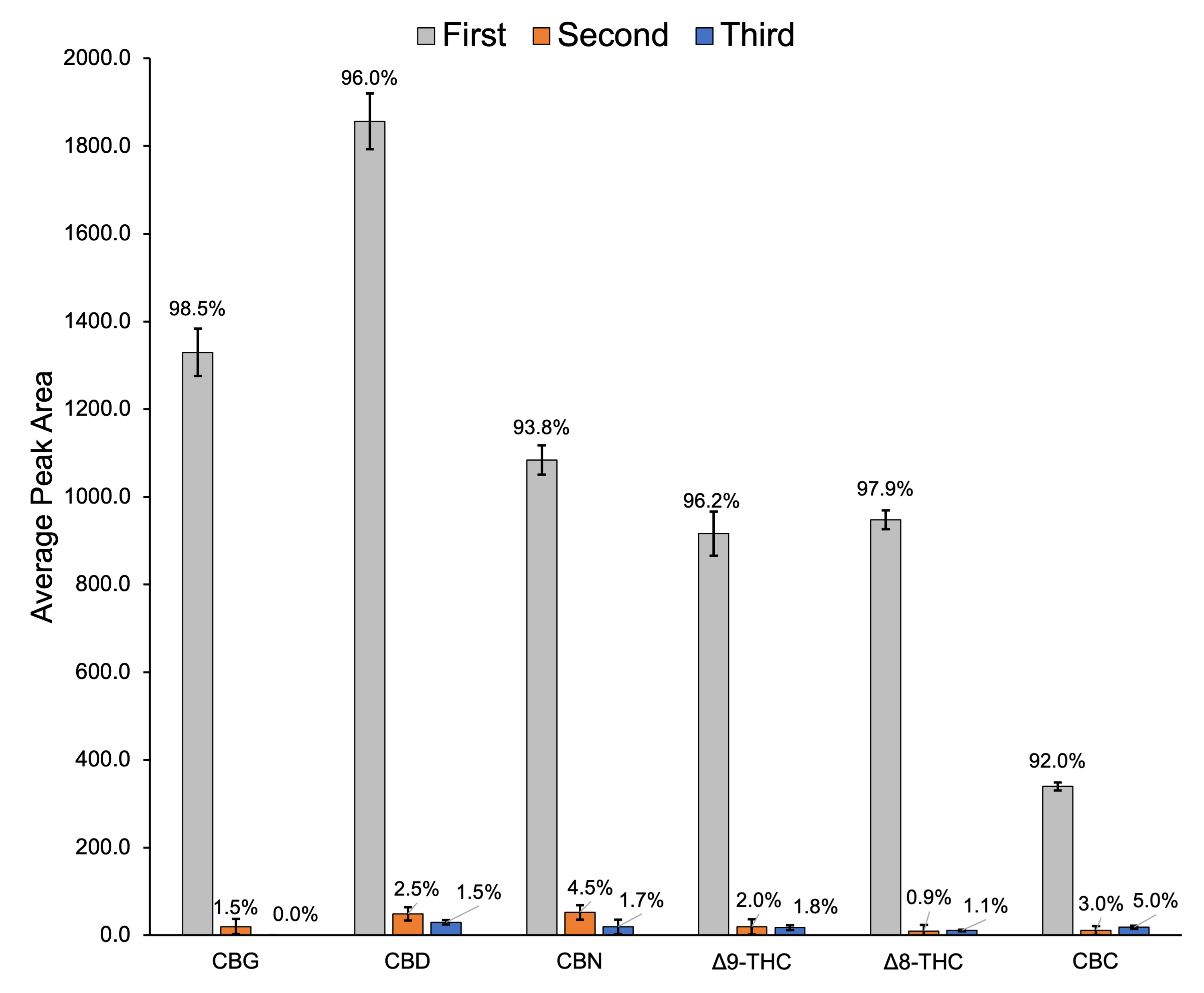

**Figure S2. Comparing relative recoveries of three subsequent extractions of 12 pooled larvae treated with 6 cannabinoids at 4 µM.** Data labels are relative percent recovery of each extraction. Error bars are SD (n=3).
